# Discovery, characterization, and application of chromosomal integration sites in the hyperthermophilic crenarchaeon *Sulfolobus islandicus*

**DOI:** 10.1101/2025.03.16.643552

**Authors:** Aashutosh Girish Boob, Changyi Zhang, Yuwei Pan, Airah Zaidi, Rachel J. Whitaker, Huimin Zhao

**Author notes:** Correspondence should be addressed to C.Z. or H.Z. These authors contributed equally to this work (Aashutosh Girish Boob, Changyi Zhang).

## Abstract

*Sulfolobus islandicus*, an emerging crenarchaeal model organism, offers unique advantages for metabolic engineering and synthetic biology applications owing to its ability to thrive in extreme environments. Although several genetic tools have been established for this organism, the lack of well-characterized chromosomal integration sites has considerably limited its potential as a cellular factory. Here, we systematically identified and characterized 13 artificial CRISPR RNAs (crRNAs) targeting eight chromosomal integration sites in *S. islandicus* using the CRISPR-COPIES pipeline and a multi-omics-informed computational workflow. By leveraging the endogenous CRISPR-Cas systems, we integrated the reporter gene *lacS* into these sites and validated heterologous gene expression through a *β*-galactosidase reporter assay, which revealed significant positional effects on expression levels. As a proof of concept, we utilized these characterized sites to genetically manipulate lipid ether composition by overexpressing GDGT (glycerol dibiphytanyl glycerol tetraether) ring synthase B (GrsB) in *S. islandicus*, a key enzyme in GDGT biosynthesis. This study expands the genetic toolbox for *S. islandicus* and highlights a concept that could be widely applicable to other *Sulfolobales*, advancing their potential as robust platforms for archaeal synthetic biology and industrial biotechnology.

## INTRODUCTION

With the increasing concerns over climate change and the unsustainable reliance on fossil fuels, there is a pressing need to transition towards more environmentally friendly and sustainable approaches for chemical production. Microorganisms play a crucial role in this transition by providing an alternate route that can help minimize the negative impacts of our current chemical industry^1^. Different microorganisms are well-suited for specific tasks^2^. For instance, anaerobic acetogens excel in the carbon-negative production of acetone and isopropanol^3^, while low-pH tolerant yeasts are ideal for organic acid production^4,5^.

Among these microbial production hosts, thermoacidophilic archaea present a particularly promising avenue^6–8^. Their distinctive attributes, including the ability to consume a wide range of substrates, thrive at high temperatures and low pH environments, and possess novel enzymes and metabolic pathways, make them attractive candidates for various bioprocesses^6^. Notably, their inherent resistance to high temperatures mitigates the risk of process failures from microbial contamination by common mesophiles^7^. Moreover, fermentation at higher temperatures improves overall kinetics, diffusion rates, and substrate solubility. In addition, it allows for easier separation of volatile products while reducing the energy costs required for fermenter cooling.

Within this context, *Sulfolobus* species hold great potential to serve as platform organisms for industrial biotechnology^9,10^. These archaea grow in high-temperature (70-85°C) and acidic (pH 2-4) environments, showcasing robust metabolic capabilities that make them well-suited for industrial applications. *Sulfolobus* species are known to consume pentose, hexose sugars, and saccharides^11^. Primarily, they have been a source of thermostable enzymes for biocatalysis and have therefore also been explored as a platform for recombinant protein production^12^. Beyond enzymes, the membranes of *Sulfolobus* species are particularly intriguing. Unlike bacteria and eukaryotes, the membranes of these archaea are composed of diether and tetraether lipids^13^, which are highly tolerant to acidic pH and high temperatures^14^. Therefore, their addition to phospholipid bilayers enhances the stability of liposomes and improves the effectiveness of oral drug delivery^15^. Furthermore, a wide variety of genetic tools, including targeted gene deletion^16–18^, inactivation^19,20^, CRISPR-Cas systems for genome editing^21,22^, and transposon insertion mutagenesis^23^, are available for these organisms. However, despite these advantages and resources, the use of *Sulfolobus* species in metabolic engineering remains largely unexplored, primarily due to their slow growth rate, limited understanding of metabolic pathways, and, most importantly, the lack of well-characterized chromosomal integration sites needed for pathway optimization.

In our study, we aim to address this gap by expanding the genetic toolkit and tapping into the unique features of a crenarchaeal model *S. islandicus*^24^. First, we employ a multi-omics approach to guide the selection of integration sites within the *Sulfolobus* genome. We then characterize these sites *in vivo* using the *lacS* (*β*-galactosidase) reporter system to ensure their suitability for gene expression. Subsequently, we leverage the characterized integration sites for strain engineering. Specifically, we modulate the lipid ether composition by overexpressing the endogenous enzyme GrsB. Overall, our work demonstrates the feasibility of engineering *S. islandicus* at highly elevated temperatures, thereby enriching the array of extreme thermoacidophiles available for biotechnological applications.

## RESULTS

### Genome-wide profiling of intergenic integration sites in *S. islandicus* M.16.4

The availability of well-characterized chromosomal integration sites plays a crucial role in understanding the unique cellular properties of an organism and in engineering strains suitable for biotechnological applications^25^. Previously, we developed CRISPR-COPIES, an easy-to-use tool for identifying intergenic loci for CRISPR-Cas-mediated gene integration^26^. The tool can discover genome-wide integration sites for bacterial and fungal genomes within minutes. We also validated the intergenic sites in *C. necator, S. cerevisiae*, and the HEK293T cell line, demonstrating its utility across diverse organisms. Therefore, we sought to assess the tool’s effectiveness in archaea, which primarily possess Type I and Type III CRISPR-Cas systems. To achieve this, we choose a hyperthermophilic archaeon, *S. islandicus* M.16.4, as a test case. This strain encodes two CRISPR-Cas subtypes: Type I-A and Type III-B Cmr-*α*^20^, which has been experimentally shown to be active against viral infections^27,28^. Type I-A recognizes the ‘CCN’ protospacer adjacent motif (PAM) located at the 5′-flanking position of the target DNA^29^, while Type III-B Cmr-*α* targets both DNA and RNA sequences^30^. However, Type III-B operates in a PAM-independent manner and relies on transcription-dependent DNA targeting^31^, rendering it less suitable for editing intergenic regions of the genome. Consequently, we opted to leverage the endogenous Type I-A CRISPR-Cas system for targeted gene insertion.

We ran the pipeline on the genome of *S. islandicus* M.16.4 to identify 40-nucleotide crRNAs that can be reprogrammed for transcription from a plasmid containing an artificial repeat-spacer-repeat array, allowing them to function within the endogenous Type I-A CRISPR-Cas system to target protospacers with a ‘CCN’ PAM and at least 10 mismatches. First, the pipeline performed a genome-wide search to locate target DNA preceded by CCA, CCT, CCG, and CCC. Next, it only selected protospacers with an 8-bp unique seed region to ensure targeting with high specificity. The pipeline utilized Scalable Nearest Neighbors (ScaNN)^32^ to filter out crRNAs that could bind to target DNA with less than 10 mismatches. As crRNAs with extreme GC content (lower than 25% or higher than 75%) tend to be less active, it filtered crRNAs falling outside this range to ensure optimal on-target activity^33^. Additionally, crRNAs containing polyN motifs (N = A, U, G, and C) were also excluded to ensure good on-target activity and efficient oligo synthesis. crRNAs targeting the protospacers with the recognition sequence of *Bsp*MI (ACCTGC) were also removed for downstream cloning experiments. Next, the pipeline procured 500 bp left and right homology arms for homologous recombination (HR)-mediated gene insertion, excluding those containing the recognition sequences of *Sal*I (GTCGAC) and *Eag*I (CGGCCG) required for cloning the donor fragment. It then selected target DNA sites in intergenic regions, ensuring a minimum distance of 250 bp from neighboring genes to prevent disruption of regulatory elements. As the bacterial or archaeal chromosomes are circular, the distal end length was set to zero, thus retaining any crRNA targeting sites situated near the annotated ends of the chromosome. Next, the pipeline utilized gene essentiality information^23^ to locate intergenic loci separated by essential genes, ensuring the stability of gene integration. The parameters used for the CRISPR-COPIES pipeline are specified in Table S1. Overall, we identified 66 crRNAs targeting 21 intergenic regions suitable for gene integration (Table S2).

It is important to note that the pipeline was previously restricted to crRNA sequences with a maximum length of 27 bp. Therefore, we updated the source code to allow searching for longer crRNAs. We also verified whether ScaNN can perform an accurate off-target search for longer crRNAs. We conducted a brute force comparison for the 66 crRNAs with all crRNAs in the genome. Only three crRNAs obtained from the pipeline had fewer than ten mismatches, indicating that ScaNN can effectively perform off-target searches for longer crRNAs (Figure S1).

Next, we integrated multi-omics information to prioritize the integration sites for characterization (Figure 1). First, we conducted a genomic analysis to determine the conservation of integration sites in *S. islandicus* strains, enabling us to identify those suitable for genetic modifications across multiple strains. We examined whether the 143-bp intergenic region (comprising a 50-nt LHR, 3-nt PAM, 40-nt protospacer, and 50-nt RHR) was conserved across the five strains of *S. islandicus*: M.16.4, REY15A (NC_017276.1), LAL14/1 (NC_021058.1), HVE10/4 (NZ_CP046616.1), and Y.G.57.14 (NC_012622.1). We sought an exact match with a 3-nt PAM and a 40-nt protospacer to identify these sites in other strains. We then used PyFAMSA^34^ to perform multiple sequence alignment. Our analysis revealed that four sites (Site 1, Site 4, Site 5, and Site 7) are well-conserved in *S. islandicus* strains, each exhibiting zero to two mutations in the 50 bp left and right homology arms (Figure S2). Subsequently, we incorporated RNA levels of neighboring genes by leveraging an in-house generated RNA-seq dataset for the RJW004 strain. This integration facilitated the identification of potentially transcriptionally active regions. Following that, we added coalescin (ClsN) information to avoid integration in repressive domains. Recent literature suggests that, like eukaryotic organisms, the archaeal chromosome organizes itself into compartments A and B, differentiated by structural maintenance of chromosomes (SMC) protein occupancy, ClsN^35^. ClsN occupancy is negatively correlated with RNA polymerase levels and transcriptional activity. Moreover, intergenic regions exhibit higher ClsN enrichment compared to open reading frames. Therefore, it is crucial to select intergenic loci with low ClsN enrichment. To achieve this, we mapped the previously described 143-bp region from M.16.4 to REY15A using BLAST^36^, obtaining mean ClsN enrichment with an alignment coverage of >90% for accurate mapping. Additionally, as the origin-containing domains (or compartment A) show higher gene expression, we computed the distances of the intergenic loci from the closest origin of replication. *S. islandicus* M.16.4 contains three origins: oriC1 (1,729,053-1,729,360), oriC2 (1-469), and oriC3 (1,261,769-1,261,941). We employed two criteria for the selection of integration sites within compartment A: mean ClsN enrichment less than two and sites located within 0.3 Mb from the nearest origin of replication. Overall, utilizing multi-omics data, we selected 13 crRNAs targeting eight integration sites for experimental validation (Table 1).

**Table 1.**
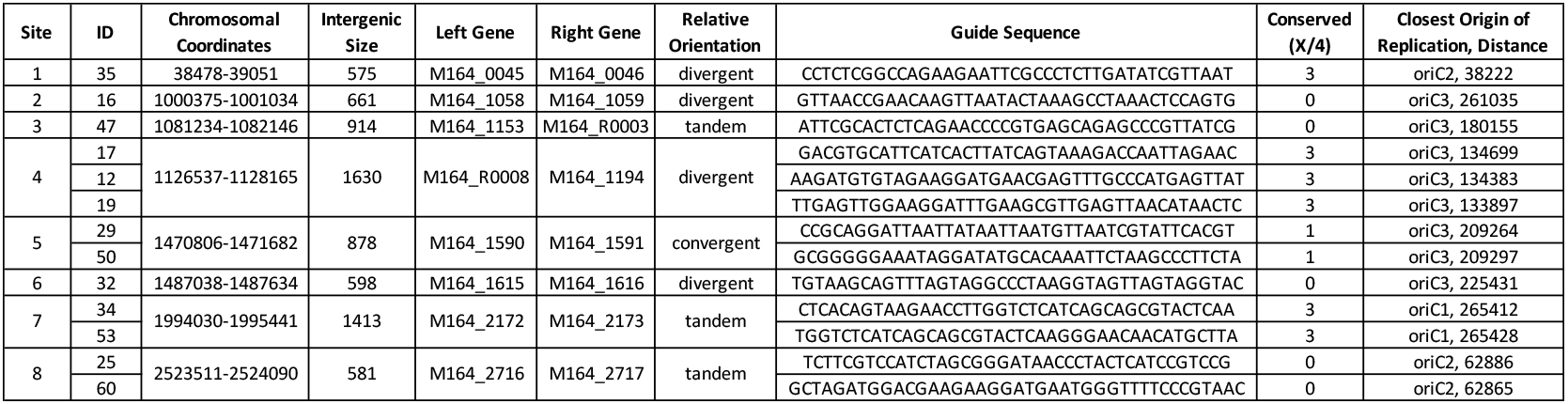
List of integration sites and the corresponding protospacers characterized in this study. We used the CRISPR-COPIES pipeline and multi-omics data integration to prioritize these sites for experimental validation.

**Figure 1.**
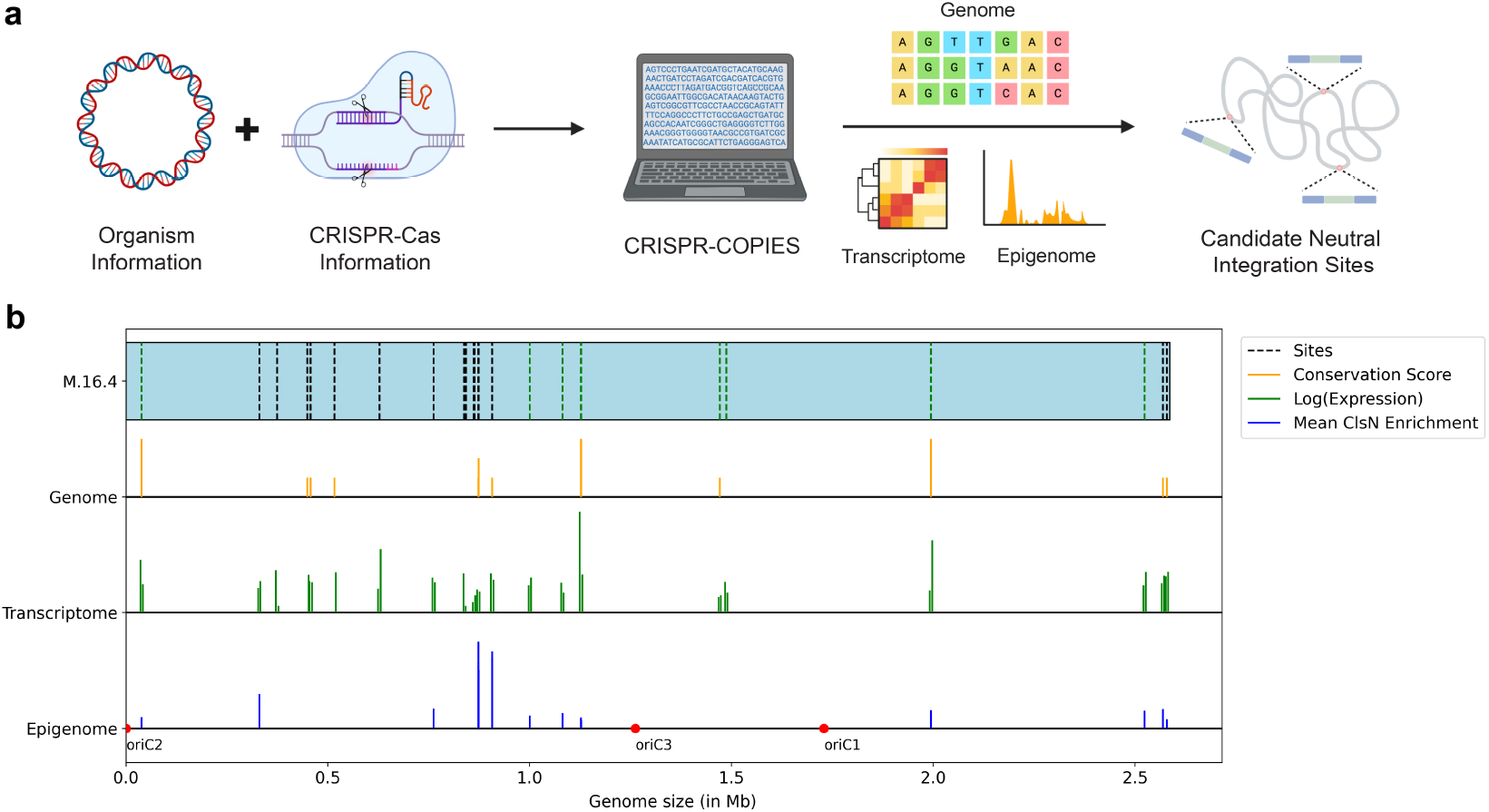
Discovery of integration sites in *S. islandicus* M.16.4. (a) Computational workflow for the selection of candidate neutral integration sites. The pipeline was executed on the genome of *S. islandicus* M.16.4 for the endogenous Type I-A CRISPR-Cas system to identify 40-nucleotide protospacers with a ‘CCN’ PAM and at least 10 mismatches. 66 crRNAs targeting 21 intergenic regions were found suitable for gene integration. Available genomics, transcriptomics, and epigenomics data were integrated to guide the selection of 13 crRNAs targeting eight intergenic loci for characterization. (b) Multi-omics map for the integration sites located using the CRISPR-COPIES pipeline. Dashed vertical lines indicate the location of the identified integration sites along the chromosome of the M.16.4 strain. The green color highlights the sites chosen for characterization in this study. The height of solid yellow, green, and blue vertical lines represents the conservation score, log_10_(expression) of neighboring genes, and mean ClsN enrichment over approximately 150 nt mapped from REY15A data using BLAST, respectively. Red circles represent the origin of replications.

### Characterization of integration sites in *S. islandicus* M.16.4

To characterize the identified sites, we constructed a series of plasmids comprising a mini-CRISPR array (repeat-spacer-repeat) and its corresponding donor DNA. The donor contained 500 bp upstream and downstream sequences serving as homology arms for integrating the *lacS* cassette from *Saccharolobus* (previously *Sulfolobus*) *solfataricus* P2 into the intergenic loci. We then transformed the plasmid into the genetic host RJW004, a derivative of M.16.4 with deletions in *pyrEF, lacS*, and *argD*. The transformants were verified by PCR analysis (Figure 2) to confirm the correct integration, followed by counterselection to cure the plasmid. While successful targeted integration was achieved at most sites, no correct colonies were obtained for site ID 60. Next, we measured the *lacS* activity to quantify the expression level from these integration sites. We obtained heterologous gene expression from all sites except ID 29 (Figure 3). As both ID 29 and ID 50 are situated within the same intergenic region, we anticipate that a mutation within the gene may account for the absence of *lacS* activity. Interestingly, our results indicated a 1.49-fold change in *lacS* activity, with the highest expression level being 1.49-fold greater than the lowest, depending on the chromosomal integration site. Moreover, we observed that plasmid-based *lacS* activity was, on average, twice as high as chromosomal activity, indicating a copy number of two for the episomal plasmid.

**Figure 2.**
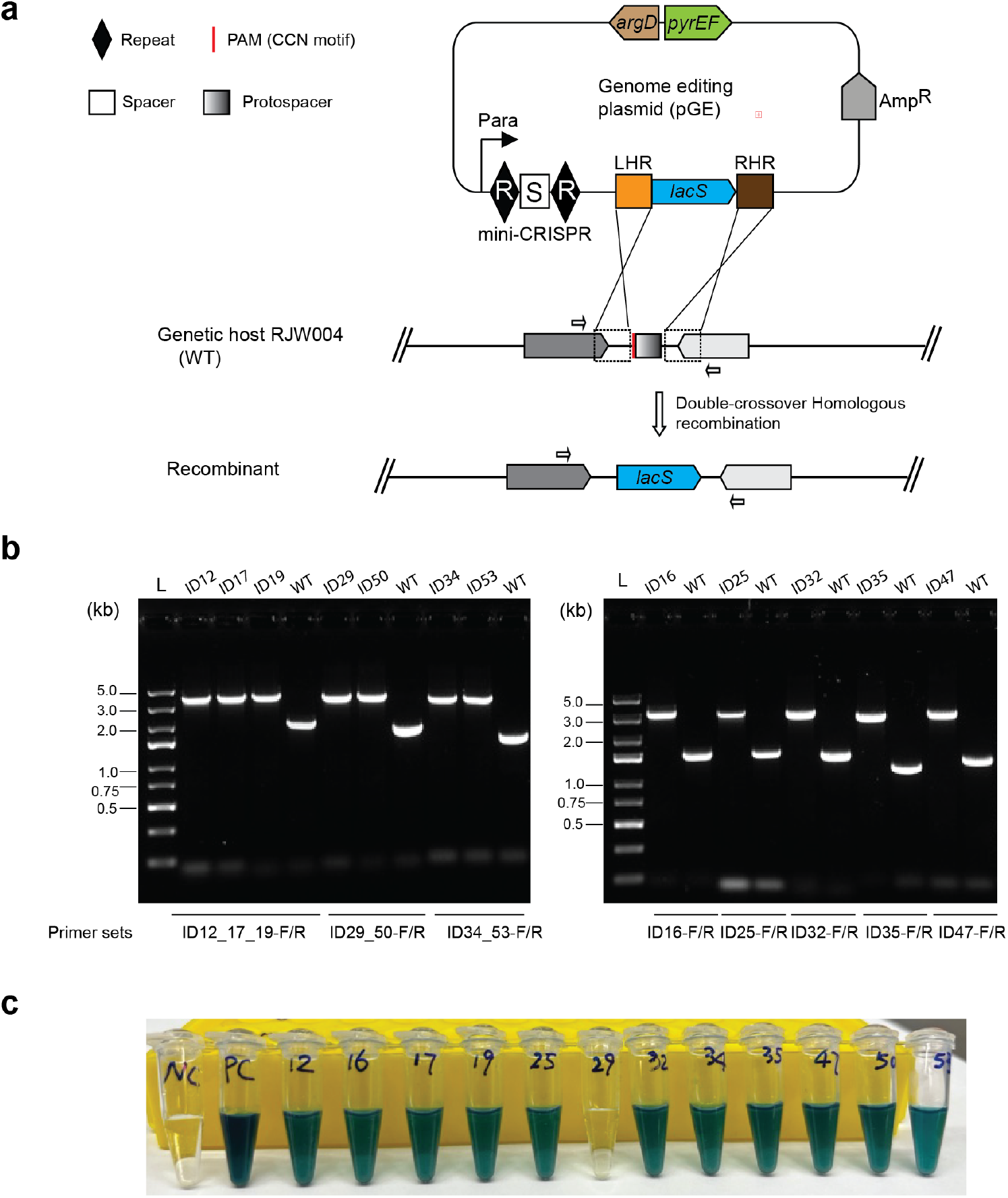
Integration of the *lacS* gene at different chromosomal sites. (a) Schematic representation of *lacS* gene integration utilizing the endogenous Type I-A CRISPR-Cas system in *S. islandicus* M.16.4. A genome editing plasmid containing a mini-CRISPR array and a donor DNA fragment with left and right homologous regions flanking the integration sites, along with the *lacS* expression cassette, was introduced into the genetic host *S. islandicus* RJW004. The Type I-A CRISPR-Cas system targeted the host chromosomal DNA, causing degradation, which was subsequently repaired through homologous recombination, leading to the generation of recombinants. Arrows indicate the relative positions of forward and reverse primers used for genotype verification. (b) PCR verification of recombinants with the *lacS* gene integrated into different chromosomal sites. Primers annealing outside the left and right homologous regions were used. The expected amplicon sizes are listed in Table S6. “L” indicates the DNA ladder, with the sizes of selected bands labeled. (c) X-gal staining of recombinants with the *lacS* gene integrated into various chromosomal sites. “NC (negative control)” refers to the RJW004 strain harboring pCYZ1, while “PC (positive control)” refers to the RJW004 strain harboring pCYZ2. Cell cultures were normalized to an OD_600_ of 0.25 and stained with X-gal (2 mg/mL) at 76°C for two hours.

**Figure 3.**
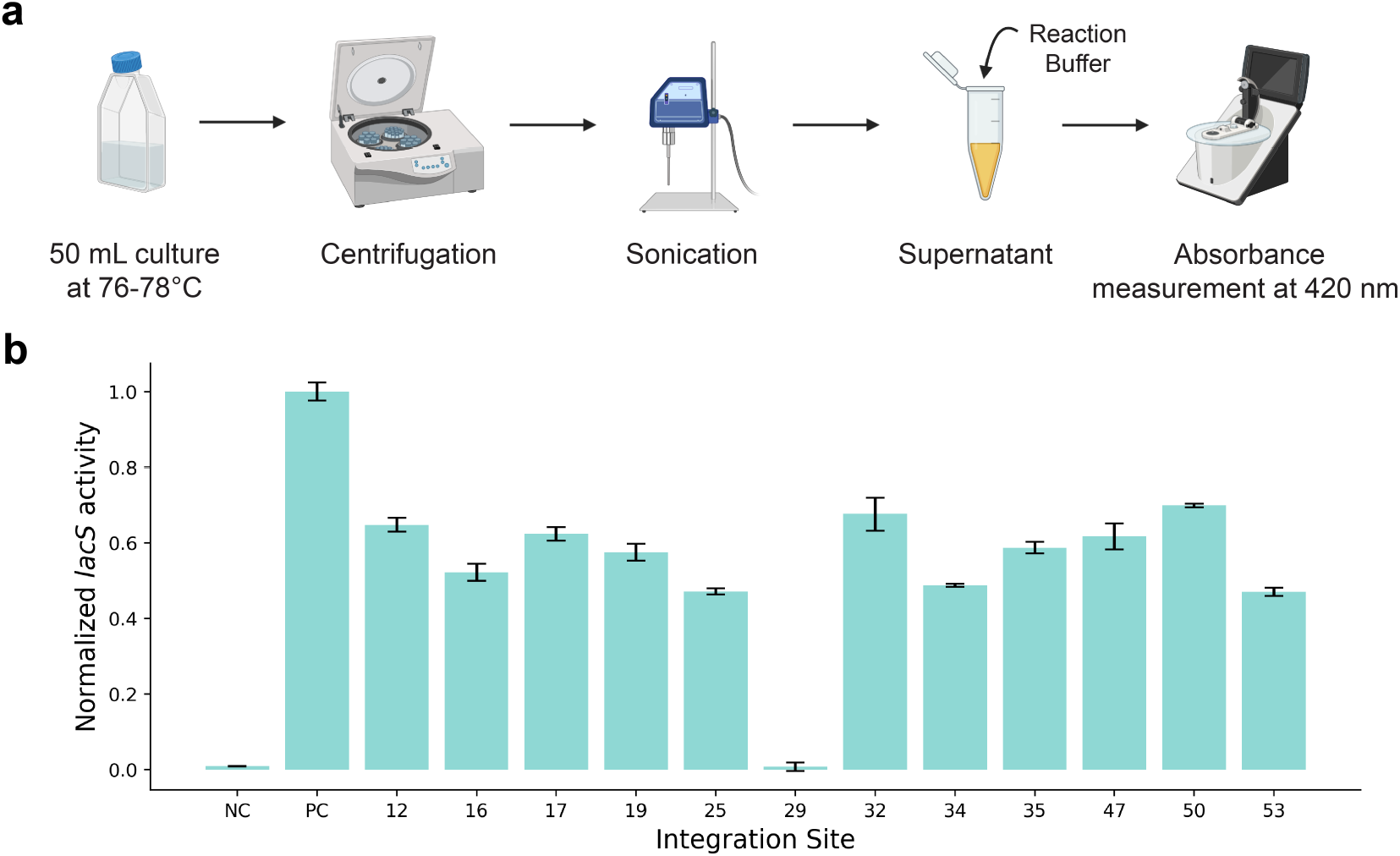
Quantification of *lacS* expression from various genomic loci using the *β*-galactosidase assay. (a) Schematic representation of the *β*-galactosidase assay workflow. Recombinant strains with the *lacS* gene integration were cultured at 76–78°C. Cell pellets were obtained via centrifugation, lysed by sonication, and the resulting supernatant was collected. The *lacS* activity was measured by adding a reaction buffer and recording the absorbance at 420 nm. (b) Normalized *lacS* activity at different chromosomal integration sites. NC: Negative Control; PC: Positive Control (plasmid-based *lacS* expression).

### Application of characterized sites for metabolic engineering

We then aimed to demonstrate the utility of the characterized sites through a metabolic engineering application focused on modulating lipid ether composition. In contrast to the ester-linked lipids predominant in bacterial and eukaryotic cell membranes, archaeal membranes are primarily composed of isoprenoid ether lipids, including glycerol diphytanyl diethers (archaeol) and glycerol dibiphytanyl glycerol tetraethers (GDGTs)^37,38^. GDGTs can contain a varying number of cyclopentane moieties, ranging from zero to eight. These GDGT cyclizations provide archaea with a survival strategy in fluctuating environmental conditions. Particularly, the number of cyclopentane moieties increases at extreme pH and temperature. These attributes render GDGTs particularly suitable for enhancing liposome stability, thereby increasing the bioavailability of drugs. Prior research has shown that liposomes containing lipids from *S. islandicus* exhibit resilience to intestinal bile salts^39^. As a result, GDGTs have found applications in oral drug delivery of therapeutic peptides^15,40,41^.

While the influence of temperature and pH on GDGT composition in archaeal membranes has been well-documented^42,43^, a recent study has elucidated the molecular mechanisms governing the expression of GDGT ring synthases (GrsA and GrsB), shedding light on their role in GDGT cyclization levels^44^. The results indicate that GrsA generates GDGT-1 to GDGT-4, followed by GrsB catalyzing the formation of GDGT-5 to GDGT-8. Notably, qRT-PCR analysis revealed that GrsB is only induced in more extreme environments, such as low pH (2.5) and high temperature (85°C). Therefore, integrating an additional copy of the *grsB* gene, driven by a robust constitutive promoter, could potentially alter GDGT composition. This modification would result in an increased number of cyclopentane moieties in lipid ethers, providing greater temperature and pH tolerance, which would be advantageous for oral drug delivery applications (Figure 4a). Hence, we amplified the endogenous *grsB* gene and cloned it into a gene expression cassette with the *Sac7d* promoter and SSV1 T6 terminator. This cassette was integrated at site ID 32 using our CRISPR-Cas-based genome editing approach (Figure 2a), followed by plasmid removal and confirmation of the strain’s genotype (Figure S3). To assess the impact of *grsB* integration, we analyzed GDGT composition in both wild-type (RJW004) and *grsB*-integrated (RJW004Int32-GrsB) strains using liquid chromatography-mass spectrometry (LC-MS). Our results showed that *grsB* integration directed lipid ether flux toward GDGTs with more cyclopentane moieties, notably GDGT-5 and GDGT-6 (Figure 4b). Specifically, a higher abundance of GDGT-5 and GDGT-6 (12.35% and 3.42% vs. 0.73% and 0.002% in wild-type) accompanied a reduction of GDGT-2 and GDGT-3 (12.02% and 20.42% vs. 14.33% and 24.45% in wild-type) in the RJW004Int32-GrsB strain, highlighting the effective use of the characterized integration site (Figure 4b and Figure S4). Moreover, chromosomal integration of the *grsB* gene did not result in any significant impact on the growth rate of RJW004Int32-GrsB (Figure 4c).

**Figure 4.**
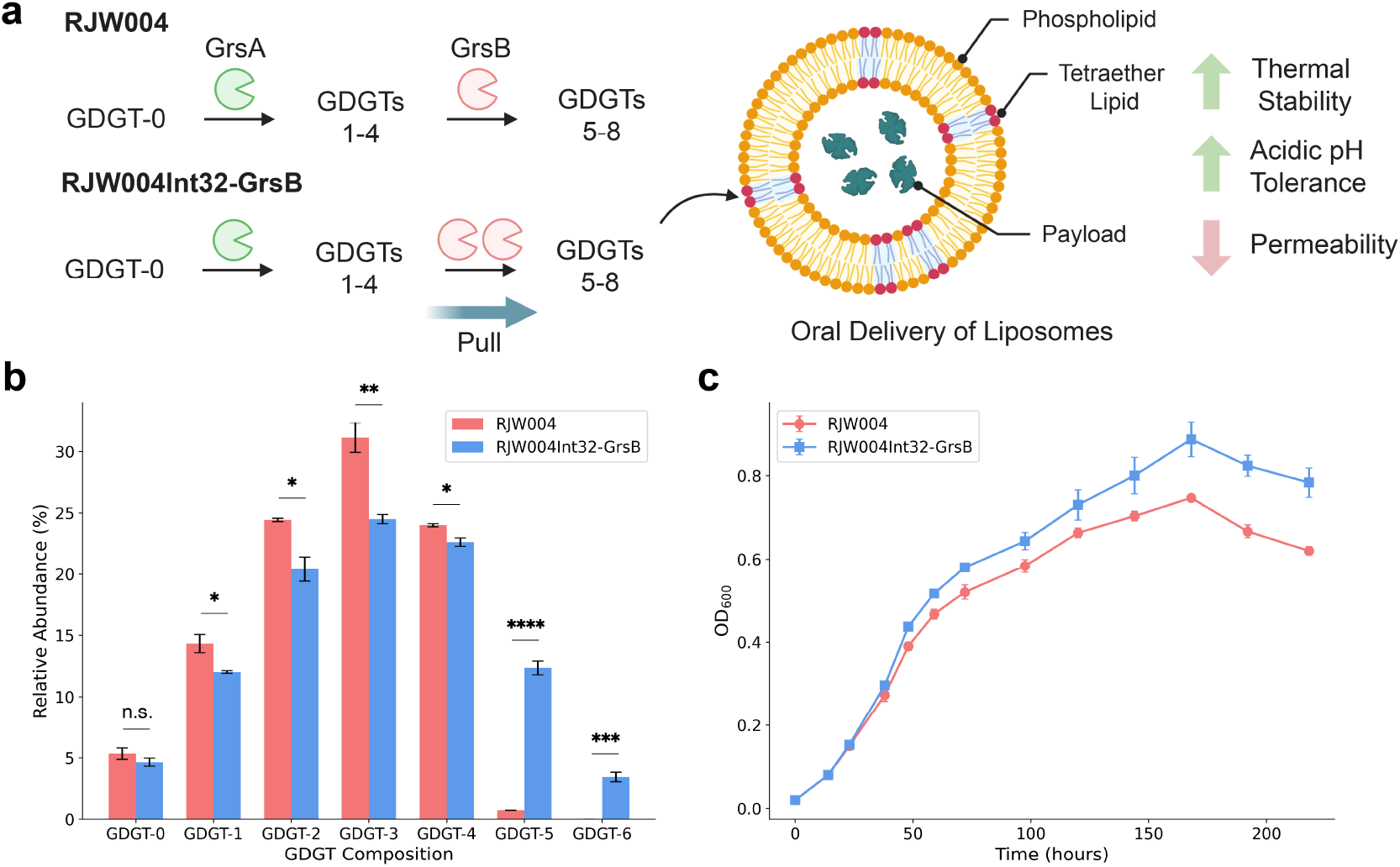
Application of the characterized integration site to modulate lipid ether composition in *Sulfolobus* strains. (a) Schematic of the GDGT cyclization pathway. An additional copy of *grsB* gene was integrated at site ID 32 to direct lipid ether flux toward more stable GDGTs with five or eight cyclopentyl rings, which could be beneficial for applications such as oral drug delivery via liposomes. (b) GDGT composition in the parental (RJW004) and the *grsB*-integrated (RJW004Int32-GrsB) strains. Integration of *grsB* resulted in enhanced production of GDGTs containing five and six cyclopentyl rings. The asterisk symbol * indicates *P* values calculated using the two-tailed unpaired *t*-test with Bonferroni correction, n.s. = not significant: *P* > 0.05, *: *P* ≤ 0.05, **: *P* ≤ 0.01, ***: *P* ≤ 0.001, ****: *P* ≤ 0.0001. (c) Growth profiles of RJW004 and RJW004Int32-GrsB strains. The strains were cultured in flasks at 76°C without shaking, and growth was monitored over nine days. Each strain was analyzed using three biological replicates. Error bars represent the standard deviation (S.D.).

## DISCUSSION

Hyperthermoacidophilic archaea are well-known as sources of thermostable enzymes. Moreover, these hosts thrive in extreme environments characterized by low pH and high temperature conditions, while obtaining energy from inorganic substrates and carbon through the CO_2_ fixation cycle^45^. Therefore, there has been increasing interest in developing them as platform strains for producing value-added biochemicals. However, despite these advantages and the availability of genetic tools, the utilization of *Sulfolobus* species in metabolic engineering remains largely untapped.

Hence in this work, we characterized integration sites that support heterologous gene or pathway expression in *S. islandicus* M.16.4, a model thermoacidophilic archaeon. We located these sites using the CRISPR-COPIES pipeline and prioritized them by integrating multi-omics datasets. Subsequently, using the *lacS* reporter system, we assayed 13 crRNAs targeting eight intergenic sites for ease of integration and gene expression. All but two crRNAs resulted in successful chromosomal integration and demonstrated *lacS* activity. Interestingly, we observed a position effect, i.e., different chromosomal locations displayed varying *lacS* activity. Finally, we demonstrated the application of our characterized sites to modulate GDGT composition. Expressing an additional copy of the *grsB* gene shifted the distribution toward more stable GDGTs containing five and six cyclopentyl rings, with minimal impact on the growth rate. These GDGTs are almost unique to extreme growth conditions, such as high temperature (85°C) or low pH (2.5), which are less conducive for *Sulfolobus* growth^46^. Therefore, our integration strategy offers a more effective approach for producing tetraether lipids with an increased number of cyclopentane moieties.

Two key challenges must be addressed to enable the production of diverse, non-native metabolites in *Sulfolobus* species. The first is its slow growth rate, which makes it less ideal as a chassis for the sustainable production of bulk chemicals. The second is the need for thermostable metabolic enzymes. Therefore, a promising direction for future development is to produce valuable fine chemicals, such as terpenes. Since archaeal membranes are made of lipid ethers derived from geranylgeranylglyceryl phosphate (GGGP), this suggests a substantial carbon flux through the isoprenoid biosynthesis pathway^47^. This presents an opportunity to use *Sulfolobus* as a host for terpene production by expressing a single heterologous enzyme. Introducing the thermostable terpene synthases (≥ 70°C), sourced from thermophilic organisms^48–50^, or engineered through ancestral reconstruction^51^ or computational design^52^, will unlock the potential of our characterized integration sites for metabolic engineering. Furthermore, these integration sites can serve as a foundation for constructing a landing pad system^53,54^, enabling multiplex engineering and pathway optimization in the future.

In conclusion, we anticipate that the integration sites identified in this study will have broad applications in both fundamental research and biotechnological advancements involving members of Sulfolobales, one of the most extensively studied groups within the TACK archaea, which share the closest common ancestor with eukaryotes after the Asgard archaea.

## MATERIAL AND METHODS

### Strains, media, and reagents

*E. coli* strain NEB10β (New England Biolabs, MA) was used for all cloning experiments. *S. islandicus* RJW004 (M.16.4 Δ*pyrEF* Δ*lacS* Δ*argD*)^55^ was used for archaeal genetic manipulation. Reagents, buffers, and growth media components were purchased from Millipore Sigma (Burlington, MA), Qiagen (Germany), or Fisher Scientific (Hampton, NH). Restriction enzymes, T4 DNA ligase, Q5 DNA polymerase, and NEBuffer 3.1 were purchased from New England Biolabs, MA. Phire Plant Direct PCR Master Mix was purchased from Thermo Fisher Scientific. All DNA oligonucleotides were ordered from Integrated DNA Technologies (Coralville, IA), while donor fragments were purchased from Twist Biosciences (San Francisco, CA). The oligos, primers, and genes used in this study are listed in Table S3.

### Plasmid construction

The backbone plasmid pCYZ1 (pSe-RP-StoargD) was generated by inserting an arginine decarboxylase expression cassette, amplified from *Sulfurisphaera tokodaii* str. 7 (StoargD), into the *Xma*I site of pSe-Rp^21^. To generate pCYZ2 (the positive control for the *β*-galactosidase activity assay), the *β*-galactosidase expression cassette, amplified from *Saccharolobus solfataricus* P2 (SsolacS), and StoargD were sequentially inserted into pCYZ1 at the *Eag*I-*Xma*I, and *Xma*I sites, respectively. Genome editing plasmids (pGE), designed for *lacS* gene integration, were created by inserting a 40 bp spacer and *lacS* donor cassette, along with the corresponding homology arms, into the pCYZ1 plasmid. In brief, to clone spacer sequences, an equimolar mixture of complementary oligos was annealed in NEBuffer 3.1. The mixture was heated to 94°C for 3 min and gradually cooled to 25°C over a period of 45 min. The annealed product was assembled into the *Paq*CI-digested pCYZ1 backbone using T4 DNA ligase. The ligation mixture was transformed in NEB-10β *E. coli* competent cells and plated on LB agar plates supplemented with appropriate antibiotics. Three colonies were picked for verification and grown overnight at 37°C in LB medium supplemented with appropriate antibiotics. The plasmid DNA was purified from the cultures using QIAprep Spin Miniprep Kit (Qiagen, Germany) following the manufacturer’s protocol. The purified plasmids were then verified using Sanger sequencing. Next, the correct plasmids were digested with restriction enzymes, *Sal*I-HF and *Not*I-HF, and assembled with 500 bp left homology arm, 500 bp right homology arm, and the *lacS* reporter gene cassette by ligation using T4 DNA ligase. In most cases, the donor fragments were obtained by digesting pTwist Amp High Copy ID plasmids with restriction enzymes, *Sal*I-HF and *Not*I-HF. For ID 17 and ID 29, left and right homology arms were PCR-amplified from the genome of *S. islandicus* using Phire Hot Start II DNA polymerase (Phire Plant Direct PCR Master Mix) and digested with *Sal*I-HF, *Fse*I and *Sbf*I, *Not*I-HF respectively to perform a four-fragment assembly with *Fse*I and *Sbf*I-digested *lacS* cassette (amplified from pCYZ2 plasmid) and *Sal*I-HF and *Not*I-HF-digested backbone plasmid pCYZ1. All constructed plasmids were verified using colony PCR, restriction digestion, and Sanger sequencing. The plasmids constructed in this study are listed in Table S4.

To construct the genome editing plasmid for integrating the *grsB* expression cassette, the plasmid pCYZ1| ID 32 was digested with *Fse*I and *Sbf*I and assembled with an empty expression cassette harboring the promoter of the *Sac7d* gene from a closely related species, *Sulfolobus acidocaldarius* DSM639^56^, and the SSV1 T6 terminator^12^. Subsequently, the constructed plasmid was digested with *Paq*CI and assembled with the *grsB* ORF amplified from the genome by ligation using T4 DNA ligase.

### *Sulfolobus* strains and growth conditions

All *S. islandicus* strains were cultivated in DT (Dextrin/EZMix N-Z-Amine A) liquid medium at 76-78°C^55^. For cultivation on plates, 2× concentrated DT medium supplemented with 20 mM MgSO_4_ and 7 mM CaCl_2_·2H_2_O was mixed in equal volumes with 1.4% (w/v) Gelrite. When needed, the medium was supplemented with 20 μg/ml uracil, 50 μg/ml agmatine and 50 μg/ml 5-Fluoroorotic Acid (5-FOA). Plates were incubated for 7–14 days in double-sealed plastic bags.

### Strain construction

The strains were constructed using S. *islandicus* RJW004, following well-established protocols as described elsewhere^20^. Plasmids were introduced via electroporation, and mutants were verified through PCR screening. Upon confirmation of successful gene integration, plasmids were eliminated using 5-FOA counterselection. All experiments were conducted under conditions previously described^20^, including medium composition, plate solidification, and growth parameters.

### *β*-Galactosidase Activity Assay

To assess the *lacS* activity, *Sulfolobus* strains were grown in 50 mL media at 76-78°C, with three biological replicates per sample. Upon reaching an OD_600_ of 0.2, cells were collected by centrifugation at 4000 rpm for 20 min and resuspended in 750 μL of 10 mM Tris-HCl, pH 8.0. Cell resuspensions were then transferred to sterile 1.5 mL microcentrifuge tubes, and samples were stored at -20°C if not processed immediately. Next, cells were lysed by sonication using a Q125 sonicator from QSonica (4 min and 30 sec with a 30-sec on, 10-sec off pulse cycle at 25% amplitude). Supernatants were then collected following centrifugation at 15,000 rpm for 10 min.

Protein concentration in the supernatant was determined using the MicroBCA Kit (Catalog number: 23235, Thermo Fisher, USA). Specifically, the supernatant was first diluted ten-fold with 10 mM Tris-HCl, pH 8.0, and the BCA reaction buffer was prepared by mixing 75 μL of Solution A, 72 μL of Solution B, and 3 μL of Solution C for each reaction. Subsequently, 150 μL of Bovine Serum Albumin (at concentrations of 0, 2.5, 5, 10, 50, and 100 ng/μL) and 150 μL of the diluted supernatant were added into a 96-well plate with lids. Following this step, 150 μL of the BCA work solution was added to each well. The plate was gently shaken for approximately 30 sec and then incubated at 37°C for two hours without shaking. After incubation, the plate was cooled to room temperature, and the absorbance at 562 nm was measured using a Tecan Infinite M1000 PRO microplate reader (Tecan Trading AG, Switzerland). A standard curve was constructed, with absorbance values from the blank well subtracted.

The *lacS* activity in the supernatant was determined using the o-nitrophenyl-β-D-galactopyranoside (ONPG) method, following a previously described protocol^57^ with slight modifications. A 10 μL of the supernatant was mixed with 490 μL of the reaction buffer (2.8 mM ONPG in 50 mM Na_2_HPO_4_-NaH_2_PO_4_ buffer, pH6.5.), incubated at 76°C for 10 min, followed by immediate termination with 500 μL of 1M Na_2_CO_3_. A control was set up by adding 10 μL of molecular biology grade water into 490 μL of reaction buffer. The absorbance of each sample at 420 nm was measured using a Nanodrop, and the concentration of ρ-nitrophenol (μmol/L) was calculated determined using a standard curve for ρ-nitrophenol.

### Quantification of protein concentration

The supernatant was first diluted ten-fold with 10 mM Tris-HCl, pH 8.0. Following this, the Bovine Serum Albumin (BCA) reaction buffer was prepared according to the MicroBCA Kit (Catalog number: 23235, ThermoFisher, USA) instructions, with 75 μL of Solution A, 72 μL of Solution B, and 3 μL of Solution C for each reaction. Subsequently, 150 μL of BCA (at concentrations of 0, 2.5, 5, 10, 50, and 100 ng/μL) and 150 μL of the diluted supernatant were added into a 96-well plate with lids. Following this step, 150 μL of the BCA work solution was added to each well. The plate was gently shaken for approximately 30 sec and then incubated at 37°C for two hours without shaking. After incubation, the plate was cooled to room temperature, and the absorbance at 562 nm was measured using a Tecan Infinite M1000 PRO microplate reader (Tecan Trading AG, Switzerland). A BCA standard curve was constructed, with absorbance values from the blank well subtracted.

### Quantification of lipid ethers in *S. islandicus* strains

To quantify GDGTs, *Sulfolobus* strains RJW004 and RJW004Int32-GrsB were grown in 50 mL of medium at 76-78°C without shaking, with three biological replicates for each sample. The cultures (50 mL) were harvested during the mid-log phase by centrifugation at 3200 × *g* for 10 min, and pellets were stored at -80°C prior to extraction. Core lipid extraction was performed following a previously reported protocol^58^. The pellets were acid-hydrolyzed with 5 mL of 10% (vol/vol) hydrochloric acid (HCl) in methanol in a capped glass vial at 70°C for 16 hours. Then, 10 mL of dichloromethane (DCM) and 10 mL of Nanopore water were added to the mixture and vortexed. The aqueous and organic phases were separated by centrifugation at 2800 × *g* for 10 min. The organic layer at the bottom was transferred to a new glass vial, and the aqueous layer was further extracted twice with 10 mL of DCM. The combined 30 mL of organic phases were dried under nitrogen to yield core lipids.

The GDGT composition was analyzed using a Q Exactive Orbitrap Mass Spectrometer (Thermo Fisher Scientific, USA) coupled with a Vanquish HPLC system (Thermo Fisher Scientific, USA). The system was equipped with a Thermo Scientific Hypersil GOLD aQ column (3 μm, 2.1 × 150 mm; PN 25303-152130) using mobile phase A (100% methanol with 0.1% formic acid) and mobile phase B (100% isopropanol with 0.1% formic acid). The column oven temperature was maintained at 60°C. GDGTs were eluted at a flow rate of 0.43 mL/min with the following gradient: an isocratic hold at 40% B for 1 min, a linear gradient from 40% B to 50% B in 9 min, an isocratic elution with 50% B for 4 min, followed by equilibration with 40% B for 1 min. MS spectra were acquired under positive ionization mode with a scan range of *m/z* 150-2000. The full scan mass spectrum resolution was set to 70,000 FWHM, the AGC target was 3 × 10^6^ ion capacity, and the maximum injection time was 100 ms. For MS/MS scans, the mass spectrum resolution was set to 17,500 FWHM, the AGC target was 1 × 10^5^ ion capacity, and the maximum injection time was 50 ms.

## ASSOCIATED CONTENT

### Supplementary Information

The Supporting Information is available free of charge on the <u>ACS Publications website</u> at DOI: XYZ.

Figures S1–S4, Table S1 (PDF)

Table S2: Genome-wide integration sites obtained for *Sulfolobus islandicus* M.16.4 using the CRISPR-COPIES pipeline with multi-omics information (XLSX)

Table S3: List of oligos, primers and genes used in this study (XLSX) Table S4: List of plasmids used in this study (XLSX)

Table S5: List of strains used in this study (XLSX)

Table S6: Expected amplicon size from different recombinants before and after the *lacS* gene integration.

## Code availability

The scripts developed in this work, intended for reproducing all analyses and regenerating figures, are available online on GitHub: https://github.com/Zhao-Group/Integration-Sites-M.16.4. The source code for the CRISPR-COPIES pipeline can be found at: https://github.com/Zhao-Group/COPIES. All scripts were developed in-house in Python 3.9. Genomes, protein sequences, and feature table for all *S. islandicus* strains were obtained from the NCBI Assembly database^59^. ClsN enrichment for *S. islandicus* REY15A was obtained from the Gene Expression Omnibus (GEO) databank using the accession code GSE128063^35^.

## AUTHOR INFORMATION

### Author contributions

A.G.B. and C.Z. conceived and designed the study. A.G.B. performed the computational experiments with input from C.Z. C.Z. developed genome editing tools based on the endogenous CRISPR-Cas systems in *S. islandicus* M.16.4 and conducted the archaeal genetic experiments. Y.P. characterized GDGTs in the archaeal strains. A.G.B., A.Z., and C.Z. constructed the plasmids. C.Z. and A.G.B. conducted *lacS* activity screening. A.G.B., C.Z., Y.P., and A.Z. analyzed the data and wrote the manuscript. H.Z. and R.J.W. secured the funding and revised the manuscript.

**Notes**

The authors declare no competing financial interest.

## ACKNOWLEDGEMENTS

This work was funded by the DOE Center for Advanced Bioenergy and Bioproducts Innovation (U.S. Department of Energy, Office of Science, Biological and Environmental Research Program under Award Number DE-SC0018420 to HZ). Any opinions, findings, and conclusions or recommendations expressed in this publication are those of the author(s) and do not necessarily reflect the views of the U.S. Department of Energy. This work was partially supported by Gordon and Betty Moore Foundation (GBMF9195 and GBMF11481 to RJW) and U.S. National Science Foundation (DBI 2400058 to HZ).

This work used the Delta system at the National Center for Supercomputing Applications through allocation BIO230077 from the Advanced Cyberinfrastructure Coordination Ecosystem: Services & Support (ACCESS) program, which is supported by National Science Foundation grants #2138259, #2138286, #2138307, #2137603, and #2138296.

## Notes

### Competing Interest Statement

The authors have declared no competing interest.

## REFERENCES

(1) Volk, M. J.; Tran, V. G.; Tan, S.-I.; Mishra, S.; Fatma, Z.; Boob, A.; Li, H.; Xue, P.; Martin, T. A.; Zhao, H. Metabolic Engineering: Methodologies and Applications. Chem. Rev. 2023, 123 (9), 5521–5570. 10.1021/acs.chemrev.2c00403.

(2) Fatma, Z.; Schultz, J. C.; Zhao, H. Recent Advances in Domesticating Non-Model Microorganisms. Biotechnology Progress 2020, 36 (5), e3008. 10.1002/btpr.3008.

(3) Liew, F. E.; Nogle, R.; Abdalla, T.; Rasor, B. J.; Canter, C.; Jensen, R. O.; Wang, L.; Strutz, J.; Chirania, P.; De Tissera, S.; Mueller, A. P.; Ruan, Z.; Gao, A.; Tran, L.; Engle, N. L.; Bromley, J. C.; Daniell, J.; Conrado, R.; Tschaplinski, T. J.; Giannone, R. J.; Hettich, R. L.; Karim, A. S.; Simpson, S. D.; Brown, S. D.; Leang, C.; Jewett, M. C.; Köpke, M. Carbon-Negative Production of Acetone and Isopropanol by Gas Fermentation at Industrial Pilot Scale. Nat Biotechnol 2022, 40 (3), 335–344. 10.1038/s41587-021-01195-w.

(4) Tran, V. G.; Mishra, S.; Bhagwat, S. S.; Shafaei, S.; Shen, Y.; Allen, J. L.; Crosly, B. A.; Tan, S.-I.; Fatma, Z.; Rabinowitz, J. D.; Guest, J. S.; Singh, V.; Zhao, H. An End-to-End Pipeline for Succinic Acid Production at an Industrially Relevant Scale Using Issatchenkia Orientalis. Nat Commun 2023, 14 (1), 6152. 10.1038/s41467-023-41616-9.

(5) Tan, S.-I.; Liu, Z.; Tran, V. G.; Martin, T. A.; Zhao, H. Issatchenkia Orientalis as a Platform Organism for Cost-Effective Production of Organic Acids. Metabolic Engineering 2025, 89, 12–21. 10.1016/j.ymben.2025.02.003.

(6) Straub, C. T.; Counts, J. A.; Nguyen, D. M. N.; Wu, C.-H.; Zeldes, B. M.; Crosby, J. R.; Conway, J. M.; Otten, J. K.; Lipscomb, G. L.; Schut, G. J.; Adams, M. W. W.; Kelly, R. M. Biotechnology of Extremely Thermophilic Archaea. FEMS Microbiology Reviews 2018, 42 (5), 543–578. 10.1093/femsre/fuy012.

(7) Zeldes, B. M.; Keller, M. W.; Loder, A. J.; Straub, C. T.; Adams, M. W. W.; Kelly, R. M. Extremely Thermophilic Microorganisms as Metabolic Engineering Platforms for Production of Fuels and Industrial Chemicals. Front. Microbiol. 2015, 6. 10.3389/fmicb.2015.01209.

(8) Ye, J.-W.; Lin, Y.-N.; Yi, X.-Q.; Yu, Z.-X.; Liu, X.; Chen, G.-Q. Synthetic Biology of Extremophiles: A New Wave of Biomanufacturing. Trends Biotechnol 2023, 41 (3), 342–357. 10.1016/j.tibtech.2022.11.010.

(9) Schocke, L.; Bräsen, C.; Siebers, B. Thermoacidophilic Sulfolobus Species as Source for Extremozymes and as Novel Archaeal Platform Organisms. Current Opinion in Biotechnology 2019, 59, 71–77. 10.1016/j.copbio.2019.02.012.

(10) Lewis, A. M.; Recalde, A.; Bräsen, C.; Counts, J. A.; Nussbaum, P.; Bost, J.; Schocke, L.; Shen, L.; Willard, D. J.; Quax, T. E. F.; Peeters, E.; Siebers, B.; Albers, S.-V.; Kelly, R. M. The Biology of Thermoacidophilic Archaea from the Order Sulfolobales. FEMS Microbiology Reviews 2021, 45 (4), fuaa063. 10.1093/femsre/fuaa063.

(11) Grogan, D. W. Phenotypic Characterization of the Archaebacterial Genus Sulfolobus: Comparison of Five Wild-Type Strains. Journal of Bacteriology 1989, 171 (12), 6710–6719. 10.1128/jb.171.12.6710-6719.1989.

(12) Peng, N.; Deng, L.; Mei, Y.; Jiang, D.; Hu, Y.; Awayez, M.; Liang, Y.; She, Q. A Synthetic Arabinose-Inducible Promoter Confers High Levels of Recombinant Protein Expression in Hyperthermophilic Archaeon Sulfolobus Islandicus. Applied and Environmental Microbiology 2012, 78 (16), 5630–5637. 10.1128/AEM.00855-12.

(13) Summons, R. E.; Welander, P. V.; Gold, D. A. Lipid Biomarkers: Molecular Tools for Illuminating the History of Microbial Life. Nat Rev Microbiol 2022, 20 (3), 174–185. 10.1038/s41579-021-00636-2.

(14) Rastädter, K.; Wurm, D. J.; Spadiut, O.; Quehenberger, J. The Cell Membrane of Sulfolobus Spp.—Homeoviscous Adaption and Biotechnological Applications. International Journal of Molecular Sciences 2020, 21 (11), 3935. 10.3390/ijms21113935.

(15) Jacobsen, A.-C.; Jensen, S. M.; Fricker, G.; Brandl, M.; Treusch, A. H. Archaeal Lipids in Oral Delivery of Therapeutic Peptides. European Journal of Pharmaceutical Sciences 2017, 108, 101– 110. 10.1016/j.ejps.2016.12.036.

(16) Deng, L.; Zhu, H.; Chen, Z.; Liang, Y. X.; She, Q. Unmarked Gene Deletion and Host–Vector System for the Hyperthermophilic Crenarchaeon Sulfolobus Islandicus. Extremophiles 2009, 13 (4), 735–746. 10.1007/s00792-009-0254-2.

(17) Wagner, M.; van Wolferen, M.; Wagner, A.; Lassak, K.; Meyer, B. H.; Reimann, J.; Albers, S.-V. Versatile Genetic Tool Box for the Crenarchaeote Sulfolobus Acidocaldarius. Front. Microbiol. 2012, 3. 10.3389/fmicb.2012.00214.

(18) Zhang, C.; Whitaker, R. J. A Broadly Applicable Gene Knockout System for the Thermoacidophilic Archaeon Sulfolobus Islandicus Based on Simvastatin Selection. Microbiology 2012, 158 (6), 1513–1522. 10.1099/mic.0.058289-0.

(19) Worthington, P.; Hoang, V.; Perez-Pomares, F.; Blum, P. Targeted Disruption of the α-Amylase Gene in the Hyperthermophilic Archaeon Sulfolobus Solfataricus. J Bacteriol 2003, 185 (2), 482– 488. 10.1128/JB.185.2.482-488.2003.

(20) Zhang, C.; Whitaker, R. J. Microhomology-Mediated High-Throughput Gene Inactivation Strategy for the Hyperthermophilic Crenarchaeon Sulfolobus Islandicus. Applied and Environmental Microbiology 2017, 84 (1), e02167–17. 10.1128/AEM.02167-17.

(21) Li, Y.; Pan, S.; Zhang, Y.; Ren, M.; Feng, M.; Peng, N.; Chen, L.; Liang, Y. X.; She, Q. Harnessing Type I and Type III CRISPR-Cas Systems for Genome Editing. Nucleic Acids Research 2016, 44 (4), e34. 10.1093/nar/gkv1044.

(22) Bost, J.; Recalde, A.; Waßmer, B.; Wagner, A.; Siebers, B.; Albers, S.-V. Application of the Endogenous CRISPR-Cas Type I-D System for Genetic Engineering in the Thermoacidophilic Archaeon Sulfolobus Acidocaldarius. Front. Microbiol. 2023, 14. 10.3389/fmicb.2023.1254891.

(23) Zhang, C.; Phillips, A. P. R.; Wipfler, R. L.; Olsen, G. J.; Whitaker, R. J. The Essential Genome of the Crenarchaeal Model Sulfolobus Islandicus. Nat Commun 2018, 9 (1), 4908. 10.1038/s41467-018-07379-4.

(24) Zhang, C.; Krause, D. J.; Whitaker, R. J. Sulfolobus Islandicus: A Model System for Evolutionary Genomics. Biochemical Society Transactions 2013, 41 (1), 458–462. 10.1042/BST20120338.

(25) Boob, A. G.; Chen, J.; Zhao, H. Enabling Pathway Design by Multiplex Experimentation and Machine Learning. Metabolic Engineering 2024, 81, 70–87. 10.1016/j.ymben.2023.11.006.

(26) Boob, A. G.; Zhu, Z.; Intasian, P.; Jain, M.; Petrov, V. A.; Lane, S. T.; Tan, S.-I.; Xun, G.; Zhao, H. CRISPR-COPIES: An in Silico Platform for Discovery of Neutral Integration Sites for CRISPR/Cas-Facilitated Gene Integration. Nucleic Acids Research 2024, 52 (6), e30. 10.1093/nar/gkae062.

(27) Athukoralage, J. S.; McMahon, S. A.; Zhang, C.; Grüschow, S.; Graham, S.; Krupovic, M.; Whitaker, R. J.; Gloster, T. M.; White, M. F. An Anti-CRISPR Viral Ring Nuclease Subverts Type III CRISPR Immunity. Nature 2020, 577 (7791), 572–575. 10.1038/s41586-019-1909-5.

(28) Bautista, M. A.; Zhang, C.; Whitaker, R. J. Virus-Induced Dormancy in the Archaeon Sulfolobus Islandicus. mBio 2015, 6 (2), 10.1128/mbio.02565-14. 10.1128/mbio.02565-14.

(29) Peng, W.; Li, H.; Hallstrøm, S.; Peng, N.; Liang, Y. X.; She, Q. Genetic Determinants of PAM-Dependent DNA Targeting and Pre-crRNA Processing in Sulfolobus Islandicus. RNA Biology 2013, 10 (5), 738–748. 10.4161/rna.23798.

(30) Han, W.; Li, Y.; Deng, L.; Feng, M.; Peng, W.; Hallstrøm, S.; Zhang, J.; Peng, N.; Liang, Y. X.; White, M. F.; She, Q. A Type III-B CRISPR-Cas Effector Complex Mediating Massive Target DNA Destruction. Nucleic Acids Research 2017, 45 (4), 1983–1993. 10.1093/nar/gkw1274.

(31) Deng, L.; Garrett, R. A.; Shah, S. A.; Peng, X.; She, Q. A Novel Interference Mechanism by a Type IIIB CRISPR-Cmr Module in Ulfolobus. Molecular Microbiology 2013, 87 (5), 1088–1099. 10.1111/mmi.12152.

(32) Guo, R.; Sun, P.; Lindgren, E.; Geng, Q.; Simcha, D.; Chern, F.; Kumar, S. Accelerating Large-Scale Inference with Anisotropic Vector Quantization. arXiv December 4, 2020. 10.48550/arXiv.1908.10396.

(33) Konstantakos, V.; Nentidis, A.; Krithara, A.; Paliouras, G. CRISPR–Cas9 gRNA Efficiency Prediction: An Overview of Predictive Tools and the Role of Deep Learning. Nucleic Acids Research 2022, 50 (7), 3616–3637. 10.1093/nar/gkac192.

(34) Deorowicz, S.; Debudaj-Grabysz, A.; Gudyś, A. FAMSA: Fast and Accurate Multiple Sequence Alignment of Huge Protein Families. Sci Rep 2016, 6 (1), 33964. 10.1038/srep33964.

(35) Takemata, N.; Samson, R. Y.; Bell, S. D. Physical and Functional Compartmentalization of Archaeal Chromosomes. Cell 2019, 179 (1), 165-179.e18. 10.1016/j.cell.2019.08.036.

(36) Camacho, C.; Coulouris, G.; Avagyan, V.; Ma, N.; Papadopoulos, J.; Bealer, K.; Madden, T. L. BLAST+: Architecture and Applications. BMC Bioinformatics 2009, 10 (1), 421. 10.1186/1471-2105-10-421.

(37) Villanueva, L.; Damsté, J. S. S.; Schouten, S. A Re-Evaluation of the Archaeal Membrane Lipid Biosynthetic Pathway. Nat Rev Microbiol 2014, 12 (6), 438–448. 10.1038/nrmicro3260.

(38) Zeng, Z.; Chen, H.; Yang, H.; Chen, Y.; Yang, W.; Feng, X.; Pei, H.; Welander, P. V. Identification of a Protein Responsible for the Synthesis of Archaeal Membrane-Spanning GDGT Lipids. Nat Commun 2022, 13 (1), 1545. 10.1038/s41467-022-29264-x.

(39) Jensen, S. M.; Christensen, C. J.; Petersen, J. M.; Treusch, A. H.; Brandl, M. Liposomes Containing Lipids from Sulfolobus Islandicus Withstand Intestinal Bile Salts: An Approach for Oral Drug Delivery? International Journal of Pharmaceutics 2015, 493 (1), 63–69. 10.1016/j.ijpharm.2015.07.026.

(40) Uhl, P.; Helm, F.; Hofhaus, G.; Brings, S.; Kaufman, C.; Leotta, K.; Urban, S.; Haberkorn, U.; Mier, W.; Fricker, G. A Liposomal Formulation for the Oral Application of the Investigational Hepatitis B Drug Myrcludex B. European Journal of Pharmaceutics and Biopharmaceutics 2016, 103, 159–166. 10.1016/j.ejpb.2016.03.031.

(41) Uhl, P.; Sauter, M.; Hertlein, T.; Witzigmann, D.; Laffleur, F.; Hofhaus, G.; Fidelj, V.; Tursch, A.; Özbek, S.; Hopke, E.; Haberkorn, U.; Bernkop-Schnürch, A.; Ohlsen, K.; Fricker, G.; Mier, W. Overcoming the Mucosal Barrier: Tetraether Lipid-Stabilized Liposomal Nanocarriers Decorated with Cell-Penetrating Peptides Enable Oral Delivery of Vancomycin. Advanced Therapeutics 2021, 4 (4), 2000247. 10.1002/adtp.202000247.

(42) Cobban, A.; Zhang, Y.; Zhou, A.; Weber, Y.; Elling, F. J.; Pearson, A.; Leavitt, W. D. Multiple Environmental Parameters Impact Lipid Cyclization in Sulfolobus Acidocaldarius. Environmental Microbiology 2020, 22 (9), 4046–4056. 10.1111/1462-2920.15194.

(43) Quehenberger, J.; Pittenauer, E.; Allmaier, G.; Spadiut, O. The Influence of the Specific Growth Rate on the Lipid Composition of Sulfolobus Acidocaldarius. Extremophiles 2020, 24 (3), 413–420. 10.1007/s00792-020-01165-1.

(44) Yang, W.; Chen, H.; Chen, Y.; Chen, A.; Feng, X.; Zhao, B.; Zheng, F.; Fang, H.; Zhang, C.; Zeng, Z. Thermophilic Archaeon Orchestrates Temporal Expression of GDGT Ring Synthases in Response to Temperature and Acidity Stress. Environmental Microbiology 2023, 25 (2), 575–587. 10.1111/1462-2920.16301.

(45) Zeldes, B. M.; Loder, A. J.; Counts, J. A.; Haque, M.; Widney, K. A.; Keller, L. M.; Albers, S.-V.; Kelly, R. M. Determinants of Sulphur Chemolithoautotrophy in the Extremely Thermoacidophilic Sulfolobales. Environmental Microbiology 2019, 21 (10), 3696–3710. 10.1111/1462-2920.14712.

(46) Jensen, S. M.; Neesgaard, V. L.; Skjoldbjerg, S. L. N.; Brandl, M.; Ejsing, C. S.; Treusch, A. H. The Effects of Temperature and Growth Phase on the Lipidomes of Sulfolobus Islandicus and Sulfolobus Tokodaii. Life 2015, 5 (3), 1539–1566. 10.3390/life5031539.

(47) Jain, S.; Caforio, A.; Driessen, A. J. M. Biosynthesis of Archaeal Membrane Ether Lipids. Front. Microbiol. 2014, 5. 10.3389/fmicb.2014.00641.

(48) Nesbitt, E. Discovering and Engineering Novel Thermostable Terpene Synthases. phd, University of Bath (United Kingdom), 2020.

(49) Lee, S.; Poulter, C. D. Cloning, Solubilization, and Characterization of Squalene Synthase from Thermosynechococcus Elongatus BP-1. Journal of Bacteriology 2008, 190 (11), 3808–3816. 10.1128/jb.01939-07.

(50) Reddy, G. K.; Leferink, N. G. H.; Umemura, M.; Ahmed, S. T.; Breitling, R.; Scrutton, N. S.; Takano, E. Exploring Novel Bacterial Terpene Synthases. PLOS ONE 2020, 15 (4), e0232220. 10.1371/journal.pone.0232220.

(51) Schriever, K.; Saenz-Mendez, P.; Rudraraju, R. S.; Hendrikse, N. M.; Hudson, E. P.; Biundo, A.; Schnell, R.; Syrén, P.-O. Engineering of Ancestors as a Tool to Elucidate Structure, Mechanism, and Specificity of Extant Terpene Cyclase. J. Am. Chem. Soc. 2021, 143 (10), 3794–3807. 10.1021/jacs.0c10214.

(52) Diaz, J. E.; Lin, C.-S.; Kunishiro, K.; Feld, B. K.; Avrantinis, S. K.; Bronson, J.; Greaves, J.; Saven, J. G.; Weiss, G. A. Computational Design and Selections for an Engineered, Thermostable Terpene Synthase. Protein Science 2011, 20 (9), 1597–1606. 10.1002/pro.691.

(53) Gaidukov, L.; Wroblewska, L.; Teague, B.; Nelson, T.; Zhang, X.; Liu, Y.; Jagtap, K.; Mamo, S.; Tseng, W. A.; Lowe, A.; Das, J.; Bandara, K.; Baijuraj, S.; Summers, N. M.; Lu, T. K.; Zhang, L.; Weiss, R. A Multi-Landing Pad DNA Integration Platform for Mammalian Cell Engineering. Nucleic Acids Research 2018, 46 (8), 4072–4086. 10.1093/nar/gky216.

(54) Fatma, Z.; Tan, S.-I.; Boob, A. G.; Zhao, H. A Landing Pad System for Multicopy Gene Integration in Issatchenkia Orientalis. Metabolic Engineering 2023, 78, 200–208. 10.1016/j.ymben.2023.06.010.

(55) Zhang, C.; Cooper, T. E.; Krause, D. J.; Whitaker, R. J. Augmenting the Genetic Toolbox for Sulfolobus Islandicus with a Stringent Positive Selectable Marker for Agmatine Prototrophy. Applied and Environmental Microbiology 2013, 79 (18), 5539–5549. 10.1128/AEM.01608-13.

(56) Berkner, S.; Wlodkowski, A.; Albers, S.-V.; Lipps, G. Inducible and Constitutive Promoters for Genetic Systems in Sulfolobus Acidocaldarius. Extremophiles 2010, 14 (3), 249–259. 10.1007/s00792-010-0304-9.

(57) Jonuscheit, M.; Martusewitsch, E.; Stedman, K. M.; Schleper, C. A Reporter Gene System for the Hyperthermophilic Archaeon Sulfolobus Solfataricus Based on a Selectable and Integrative Shuttle Vector. Molecular Microbiology 2003, 48 (5), 1241–1252. 10.1046/j.1365-2958.2003.03509.x.

(58) Zeng, Z.; Liu, X.-L.; Farley, K. R.; Wei, J. H.; Metcalf, W. W.; Summons, R. E.; Welander, P. V. GDGT Cyclization Proteins Identify the Dominant Archaeal Sources of Tetraether Lipids in the Ocean. Proceedings of the National Academy of Sciences 2019, 116 (45), 22505–22511. 10.1073/pnas.1909306116.

(59) Sayers, E. W.; Bolton, E. E.; Brister, J. R.; Canese, K.; Chan, J.; Comeau, D. C.; Connor, R.; Funk, K.; Kelly, C.; Kim, S.; Madej, T.; Marchler-Bauer, A.; Lanczycki, C.; Lathrop, S.; Lu, Z.; Thibaud-Nissen, F.; Murphy, T.; Phan, L.; Skripchenko, Y.; Tse, T.; Wang, J.; Williams, R.; Trawick, B. W.; Pruitt, K. D.; Sherry, S. T. Database Resources of the National Center for Biotechnology Information. Nucleic Acids Research 2022, 50 (D1), D20–D26. 10.1093/nar/gkab1112.

